# Assay for characterizing adsorption-properties of surfaces (APS) used for sample preparation prior to quantitative omics

**DOI:** 10.1101/2023.08.03.551632

**Authors:** Bente Siebels, Manuela Moritz, Diana Hübler, Antonia Gocke, Hannah Voß, Hartmut Schlüter

**Author notes:** **Corresponding Author** Hartmut Schlüter, Section / Core Facility Mass Spectrometric Proteomics, Diagnostic Center, University Medical Center Hamburg–Eppendorf, Martinistraße 52, 20251 Hamburg, Germany.

## Abstract

Analytes during their journey from their natural sources to their identification and quantification are prone to adsorption to surfaces before they enter an analytical instrument, causing false quantities. This problem is especially severe in diverse omics. Here, thousands of analytes with a broad range of chemical properties and thus different affinities to surfaces are quantified within a single analytical run. For quantifying adsorption effects caused by surfaces of sample handling tools, an assay was developed, applying LC-MS/MS-based differential bottom-up proteomics and as probe a reference mixture of thousands of tryptic peptides, covering a broad range of chemical properties. The assay was tested by investigating the adsorption properties of several vials composed of polypropylene, including low-protein-binding polypropylene vials, borosilicate glass vials and low-retention glass vials. In total 3531 different peptides were identified and quantified across all samples and therefore used as probes. A significant number of hydrophobic peptides adsorbed on polypropylene vials. In contrast, only very few peptides adsorbed to low-protein-binding polypropylene vials. The highest number of peptides adsorbed to glass vials, driven by electrostatic as well as hydrophobic interactions. Calculation of the impact of the adsorption of peptides on differential quantitative proteomics showed significant false results. In summary, the new assay is suitable to characterize adsorption properties of surfaces getting into contact with analytes during sample preparation, thereby giving the opportunity to find parameters for minimizing false quantities.

**Insert Table of Contents artwork here:** 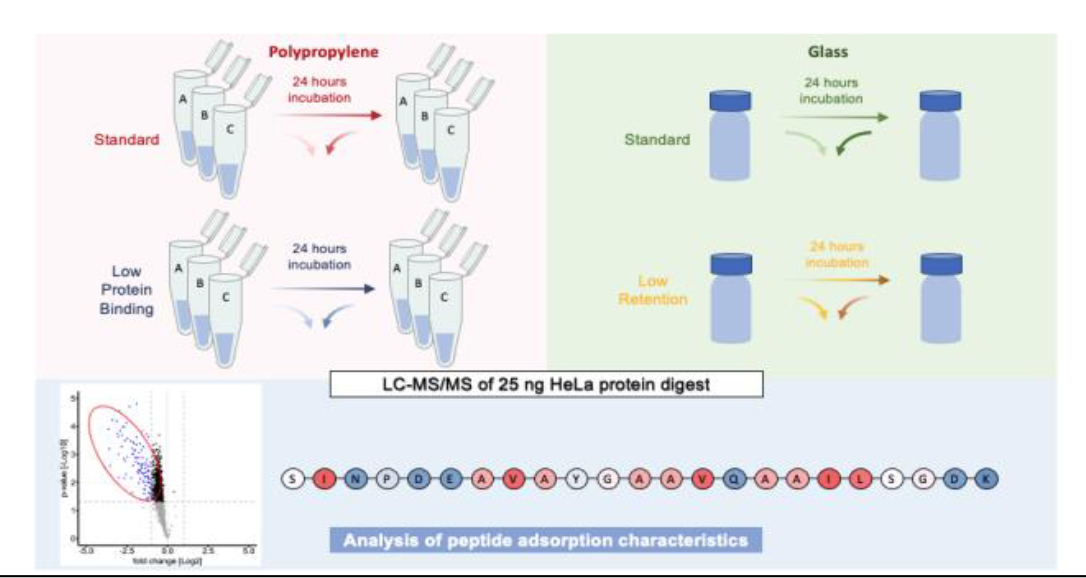

## INTRODUCTION

For quantification many analytical instruments are requiring analytes in a solubilized form. If analytes are not already dissolved in liquids like body fluids, they must be transferred from their original matrix into a sample solvent. Tissues usually are homogenized for releasing the analytes. Thereafter, additional sample preparation steps such as removal of interfering molecules or enrichment of analytes may be required. During the journey from their natural source to the quantification instrument, the analytes pass diverse vessels and tubing, often consisting of different chemistries. If analytes have a high affinity towards molecules of the surfaces they will be adsorbed resulting in the decrease of the total amount of analytes and consequently, finally false-negative results will be obtained. If only a single or few analytes will be quantified their loss during sample preparation can be compensated by the application of internal standards. However, in omics studies usually thousands of analytes are quantitated. For such studies the integration of internal standards for each of the thousands of analytes is too expensive. Furthermore, in many omics studies the identities of the analytes are not known and thus internal standards not applicable.

Currently, new developments in omics are focusing on improving sensitivity for enabling single cell omics^1^ and even sub-cellular omics^2^. Technical advances in the development of mass spectrometer systems enable bottom-up proteome analysis with total protein amounts less than one ng.^3^ Vials, used in omics are commonly made of polypropylene or borosilicate glass, because they are inexpensive and compatible with most solvents. Since the vials from different vendors are different according to their chemistry, the affinities of their surfaces towards sample molecules differ also, which was described with respect to adsorption of molecules such as pharmaceutics^4^, peptides or proteins^5^. Borosilicate glass is composed of SiO_2_ and has, even under very acidic conditions, a negatively charged surface, attracting positively charged molecules.^6^ In contrast, polypropylene is uncharged and attracts hydrophobic molecules.^7^ In consequence, adsorption can cause loss of analytes, leading to inaccurate quantification and identification of biomolecules^8^.

To reduce sample loss in omics “anti-adsorption diluents”^4,9^ and surfactants, such as n-dodecyl-β-D-maltoside (DDM)^10^ are frequently used. However, these substances can interfere with the ionization and desorption of biomolecules during mass spectrometric analysis ^10^, increase sample complexity and background noise^11^ and thereby lead to the distortion of quantitative values. An alternative is the use of chemically modified surfaces, minimizing adsorption of analytes to vials.

Especially proteins are known to adsorb to surfaces because they have many different functional groups with different chemical properties. Thus, adsorption of various proteins to surfaces, has been widely characterized.^12,13^ However, a major problem of studies using proteins is, that the exact composition of atoms of proteins most often is not known, especially with respect to their posttranslational modifications. Furthermore, usually a protein coded by a single specific gene is not present as a single molecule but as a mixture of different proteoforms^14,15^, often showing very different chemical properties. In contrast to proteins the chemical properties of peptides are clearly defined by their amino acid sequences. Therefore, peptides are better suited than proteins for studying which chemical properties of molecules will favor adsorption to surfaces. In addition, peptides have a less complex 3-dimensional, more linear structure than proteins, which may hide hydrophobic amino acids on the inside by folding, thereby increasing the problem that the calculated hydrophobicity is not correlating with the observed hydrophobicity^16^. Surprisingly, only few studies focus on the investigation of adsorption phenomena of molecules on surfaces using peptides as model compounds. Most of these studies used single synthetic peptides or low complexity protein digests^17^. The characterization of adsorption of complex peptide mixtures, typically present in tryptic digest of protein extracts of cells, tissues, or body fluids, is still missing, although such an approach is associated with the advantage, that in a complex mixture of tryptic peptides many peptides with very different chemical properties are present.

The loss of molecules during sample handling is significantly increasing with their decreasing concentration caused by their adsorption towards surfaces like reaction vials. Therefore, it is important to find surfaces with minimal affinity towards the analytes. We developed a method for quantifying loss of individual analytes with different chemical properties by using a complex peptide mixture, which is a kind of extension of the “DMD-test” mixture used for characterizing retention behavior of reversed phase columns presented by Daldrup et al in 1984^18^. We targeted the question how well the chosen complex reference peptide mixture, which represents the main idea of the assay for characterizing adsorption-properties of surfaces (APS), is suitable for investigating the adsorption characteristics of surfaces with different chemical compositions, getting into contact with analytes, dissolved in aqueous liquids. The applicability of APS was tested by answering the question, if surfaces of vials consisting of different chemistries, have a significant impact on the quantitative bottom-up proteomics results.

## EXPERIMENTAL SECTION

### Sample preparation

HeLa protein-digest standard (Pierce, Thermo Fisher Scientific, Schwerte, Germany) was dissolved in 0.1% formic acid (FA) to a concentration of 0.25 μg/μL. Polypropylene standard safe-lock microcentrifuge vials and corresponding low protein binding vials of 1.5 mL size from three manufacturers were used: Manufacturer A (Eppendorf Safe-Lock Tubes no. 0030120086 and Protein LoBind® Tubes no. 0030108116, Eppendorf SE, Hamburg, Germany), Manufacturer B (Pierce™ Microcentrifuge Tubes no. 69715 and Low Protein Binding Microcentrifuge Tubes no. 90410, Thermo Fisher Scientific, Life Technologies GmbH, Carlsbad, USA) and manufacturer C (SafeSeal reaction tube no. 72.706 and SafeSeal reaction tube, Low protein-binding no. 72.706.600, Sarstedt AG & Co. KG, Nümbrecht, Germany). Further, standard 1 ml glass vials (1 mL LCGC Certified Clear Glass, Total Recovery, Waters, Milford, USA) and respective low retention glass vials (1 mL TruView pH Control LCMS Certified Clear Glass, Total Recovery, Waters, Milford, USA) were used.

Directly prior to the measurement HeLa protein-digest standard was dissolved to a final concentration of 5 ng/μL (total volume 100 μL), with 0.1 % FA in each vial. For polypropylene vials 10 μL were immediately transferred into autosampler glass vials. Without delay 25 ng (5μL) HeLa protein-digest standard control samples were immediately injected from the autosampler vial into the LC-MS system. (Control, 0h).

To measure peptide adsorption effects of the surfaces of the vials, HeLa peptides were incubated in the respective vials for 24 hours at room temperature. After 24h incubation, 10 μL of the samples were transferred to autosampler glass vials 5μL were immediately injected into the LC-MS system. For glass, 5 μL were directly injected from the glass vials incubated in the autosampler at 4°C.

### LC-MS/MS acquisition

LC-MS analysis was performed on a nano UPLC (nanoAcquity system, Waters, Milford, USA), coupled to a quadrupole orbitrap hybrid mass spectrometer (QExactive, Thermo Fisher Scientific, Waltham, USA). Chromatographic separation of peptides was achieved with a two-buffer system (buffer A: 0.1% FA in water, buffer B: 0.1% FA in ACN). For online desalting and purification, a peptide trap (180 μm × 20 mm, 100 Å pore size, 5 μm particle size, Symmetry C18, Waters) was installed in front of a 25 cm C18 reversed phase column (75 μm × 200 mm, 130 Å pore size, 1.7 μm particle size, Peptide BEH C18, Waters). Elution of the peptides occurred with an 80 min gradient with linearly increasing concentration of buffer B from 2% to 30% in 60 min, rising to 90% for 5 min with equilibration for 10 min at 2% buffer B. Eluted peptides were ionized and desorbed via electrospray ionization, using a spray voltage of 1.8 kV. The ions being responsible for the 15 highest signal intensities per precursor scan (1 × 106 ions, 70,000 Resolution, 240ms fill time) were analyzed by MS/MS (HCD at 25 normalized collision energy, 1 × 105 ions, 17,500 Resolution, 50 ms fill time) in a range of 400–1200 m/z. A dynamic precursor exclusion of 20 s was used.

### Database searching

Acquired spectra from LC-MS/MS measurements were processed in Proteome Discoverer software (Version 2.41.15), Thermo Fisher Scientific, Milford, USA) and searched against a reviewed human Swissprot database, obtained in April 2021 containing 20 365 entries. As fixed modifications carbamidomethylation of cysteine residues was set. Pyroglutamate formation at glutamine residues, acetylation and methionine loss were set as dynamic modifications of the N terminus. Oxidation of methionine and formylation of serine, lysine and threonine were set as dynamic modifications. A maximum number of 2 missing tryptic cleavages was set. Only peptides between 6 and 144 amino acids where considered. A strict cut-off (FDR<0.01) was set for peptide identification. Quantification was performed using the Minora Algorithm implemented in Proteome discoverer.

### Statistical analysis and visualization at the peptide level

To measure adsorption effects, areas under the curves of the extracted-ion chromatograms of identified peptides, were loaded into the statistical analysis program Perseus (Version 1.5.8.5., Max-Planck Institute of Biochemistry, MaxQuant, Munich, Germany)^19^.

Peptide abundances were log2 transformed and used for further analyzes. No normalization was applied. Linear principal components analysis (PCA) was carried out using complete observations, to visualize similarities and differences between samples. The first three principal components (explained variance > 5%) were considered. In addition, Pearson correlation coefficients were calculated as similarity metrics between individual samples within and between different setups. In boxplots, 50% of the data points are inside the box (Q1 (Quartile 1) being the lower bound of the box (25%), Q3 being the upper bound of the box (75%)). Whiskers show all values beyond the box without outliers. Outliners were defined as Q3 + 1.5 * IQR (Interquartile range) (upper outlier) and Q1-1.5 * IQR (lower outlier). IQR being Q1–Q3.

For further analysis, only peptides identified across all setups were used to compare adsorption effects between different polypropylene vials and glass vials. Oxidized, formylated and pyroglutamate peptides as well as their unmodified counterpart were removed, as their modification was traced back to the incubation in 0.1% formic acid^20-22^.

To identify significantly adsorbed peptides Welch’s t-testing was performed comparing peptide abundancies of the control sample (0h) and sample incubated 24h for all surfaces. Peptides with a p-value ≤ 0.05 and a fold-change (FC) ≥ 2-fold between the compared groups were considered as statistically significant differential abundant. Respective peptides were classified as adsorbed in the individual setup. T-testing results were visualized in a volcano plot, plotting the - log10(p-value) against the log2 fold-change, using the APS test in house script developed in Python (Version 3.10.9) using the Spyder IDE (Version 5.4.1) software environment.

To compare adsorbed peptides between different containers the package “UpSetR”^23^ was used in the R software environment^24^.

### Analysis of chemical properties of the adsorbed peptides

For the characterization of chemical properties of adsorbed peptides, the APS test in house script was used developed in Python (Version 3.10.9) using the Spyder IDE (Version 5.4.1) software environment.

Figures in this manuscript were created using GraphPad Prism (Version 8.0.2, San Diego, California USA)^25^. Charge states below 2+ were neglected in the analysis since only charge states from 2-6 were included during MS-acquisition. To detect significant differences between peptide properties of adsorbed and non-binding peptides a two-sided Wilcoxon rank sum test with continuity correction was performed in the R software environment^24^.

For comparing the distribution of amino acids in adsorbed and non-binding peptides, the peptide of 15 amino acids in length being most commonly adsorbed was selected in Microsoft Excel^26^. Values were calculated with an in-house R script based on the individual amino acid proportion within 15 AAs long peptides and percentage in adsorbed peptides was subtracted from non-binding peptides.

### Protein quantification and analysis

To measure the effect of peptide adsorption on protein quantification, peptide abundances from unique peptides identified in all studies, were summed with the consolidate function in Microsoft Excel according to their protein of origin. Respective abundances were log2 transformed. Welch’s T-testing was performed between control sample (0h) and sample incubated 24h for each setup. Proteins identified with a p-value ≤ 0.05 and a fold-change ≥ 1.5 or ≥ 2 were considered as significantly differential abundant. T-testing results were visualized in a volcano plot, plotting the -log10(p-value) against the log2 fold-change, using a in house script in the R software environment^27^.

## RESULTS AND DISCUSSION

### Testing the assay for characterizing adsorption-properties of surfaces (APS)

For the applicability test of APS for measuring the adsorption characteristics of surfaces diverse vials from different manufacturers, comprising polypropylene (PP) vials, polypropylene-based low-protein-binding (LPB) vials, as well as glass vials (G) (unmodified borosilicate glass) and low-retention (LR) glass vials were chosen. The vials were incubated 24 hours with a commercially available reference tryptic peptide mixture derived from HeLa cells, commonly used for quality control. Relative peptide quantities of the control sample (injection of the tryptic peptides directly after dissolving the peptides, termed 0h) and the sample incubated 24h was assessed by differential quantitative proteomics using label free quantitative liquid-chromatography-tandem mass spectrometry (LC-MS/MS). In total 3531 individual peptides were identified and quantified across all samples (Supplementary Table S1-2).

Principal component analysis (PCA) revealed significant adsorption mechanisms in PP vials, visible as clear differentiation based on principal components 1 and 2 of all PP vial replicates, after 24 hours incubation (Figure S1. a.). For LPB, significant differences between the control sample (0h) and sample incubated 24h were detectable for manufacturer B. Significant adsorption of peptides was also detected after incubation of the peptides in LR and G vials. Between replicates of each PP-based vial a Pearson correlation > 96 % was observed, for glass vials a Pearson correlation > 98 % was observable for the control samples (Figure S1. b.) and proofs the reproducibility of the developed assay.

To identify peptides that adsorbed to PP and G from the 3531 quantified peptides, Welch’s T-testing (p-value ≤ 0.05) was performed, comparing the peptide abundances between the control sample and sample incubated 24h for each vial. Peptides, that showed a p-value significance and 2-times lower abundance after 24h-incubation were classified as *adsorbed* (Figure 1a).

**Figure 1.**
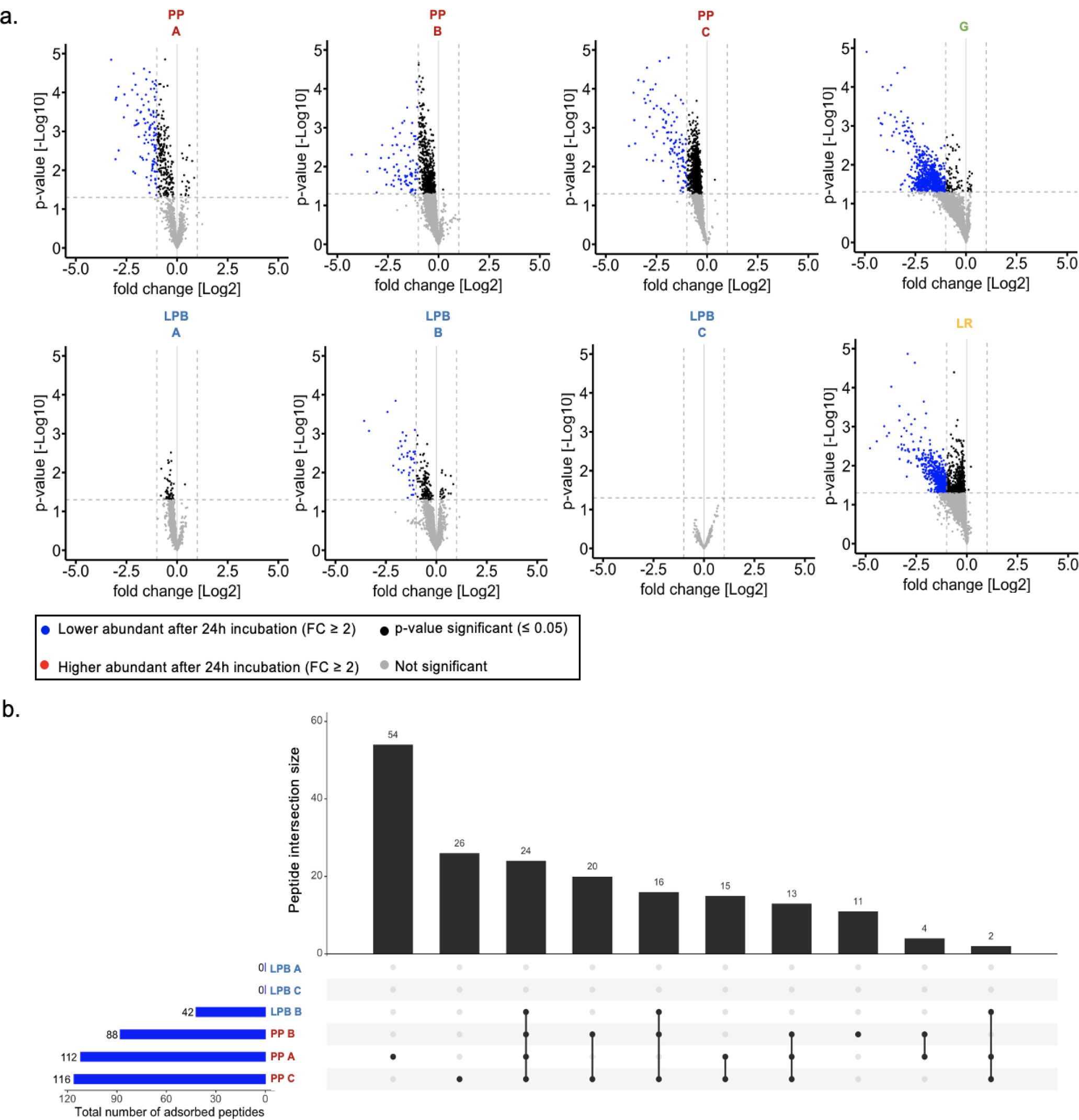
Adsorbed peptides after 24 hours incubation in polypropylene (PP) vials or glass vials. a. Overview about number of significantly adsorbed peptides. Volcano plots show the –log10 p-value against the log2 foldchange difference for the comparison between the control sample (0h) and the sample incubated 24h, individually determined for low protein binding (LPB) polypropylene vials, PP vials, glass vials (G) and low retention glass vials (LR). For APS only those peptides (N=3531) were used which were detected across all samples. Formylated, oxidised and pyro-glutamated peptides and their unmodified counteract were removed. Peptides identified with a p-value ≤ 0.05 and a fold-change difference ≥ 2 were considered significantly adsorbed. APS test results can be found in supplementary Table S3. b. Upset Venn diagram showing the overlap of adsorbed (p-value ≤ 0.05, fold-change (FC) difference ≥ 2) peptides after 24 hours incubation in PP and LPB vials from manufacturers A, B and C. Blue bar diagram represents the total number of adsorbed peptides per PP-vial. Black dots in the matrix indicate the peptide overlap between different vials with intersection size shown in the black bar diagram above.

In our study we observed that unmodified polypropylene vials adsorbed significant numbers of peptides, when stored in 0.1% formic acid (FA) in water, commonly used in LC-MS/MS experiments in proteomics. For PP vials the highest number of adsorbed peptides after 24h was detected in vials from manufacturer C (A: 112, B: 88, C: 116). In contrast, LPB vials of manufacturers A and C did not show significant peptide adsorption. The LPB vials of manufacturer B significantly adsorbed 42 peptides. An even more significant effect, with respect to peptide adsorption, compared to polypropylene was observed for G and LR vials. Here, more than 18% of all identified peptides significantly adsorbed to the glass vials (G: 812 adsorbed peptides; LR: 655 adsorbed peptides). These results demonstrate that with APS subtle differences between chemically similar surfaces are detectable. Furthermore, APS showed that the surface modification of polypropylene vials was effective in significantly reducing peptides adsorbing to the surface of vials.

Comparing PP vials between all manufacturers, 24 (10.6%) of 227 peptides adsorbed to PP vials from all manufacturers (Figure 1b). 54 peptides only adsorbed to PP vials of manufacturer A, while 26 peptides only adsorbed to PP vials from manufacturer C. No peptide was only adsorbed to PP from manufacturer B. Peptides, adsorbed to LPB from manufacturer B also adsorbed to PP vials. 593 (67.8 %) of the adsorbed peptides bound to both, G vials and LR vials. 62 peptides exclusively adsorbed to LR vials, while 219 peptides exclusively adsorbed to G vials (Figure S2). While the exact chemistry of modified LPB across different manufacturers is not available, it can be assumed, that these differences between different manufacturers can be traced back to different methods for PP modification and PP manufacturing processes in order to make PP surfaces less hydrophobic and smoother^28^, or in the case of LR vials less ionic. As a high overlap between peptides adsorbed to LR and G, as well as LPB from manufacturer B and PP was detected, only peptide properties from peptides, adsorbed to PP and G surfaces were further analyzed.

### APS assay enables for the chemical characterization of adsorbed peptides

To investigate the chemical peptide properties leading to adsorption to PP and G vials, peptide length, hydrophobicity (GRAVY number), and peptide charge at pH = 2.7 (pH-value of 0.1% FA) were calculated for adsorbed and non-binding peptides for each vial type (Figure 2). Subsequently, results were compared to chemical interactions known from the literature to validate the APS assay quantitative results and enable even more specific characterization of analyte and surface on the amino acid level.

**Figure 2.**
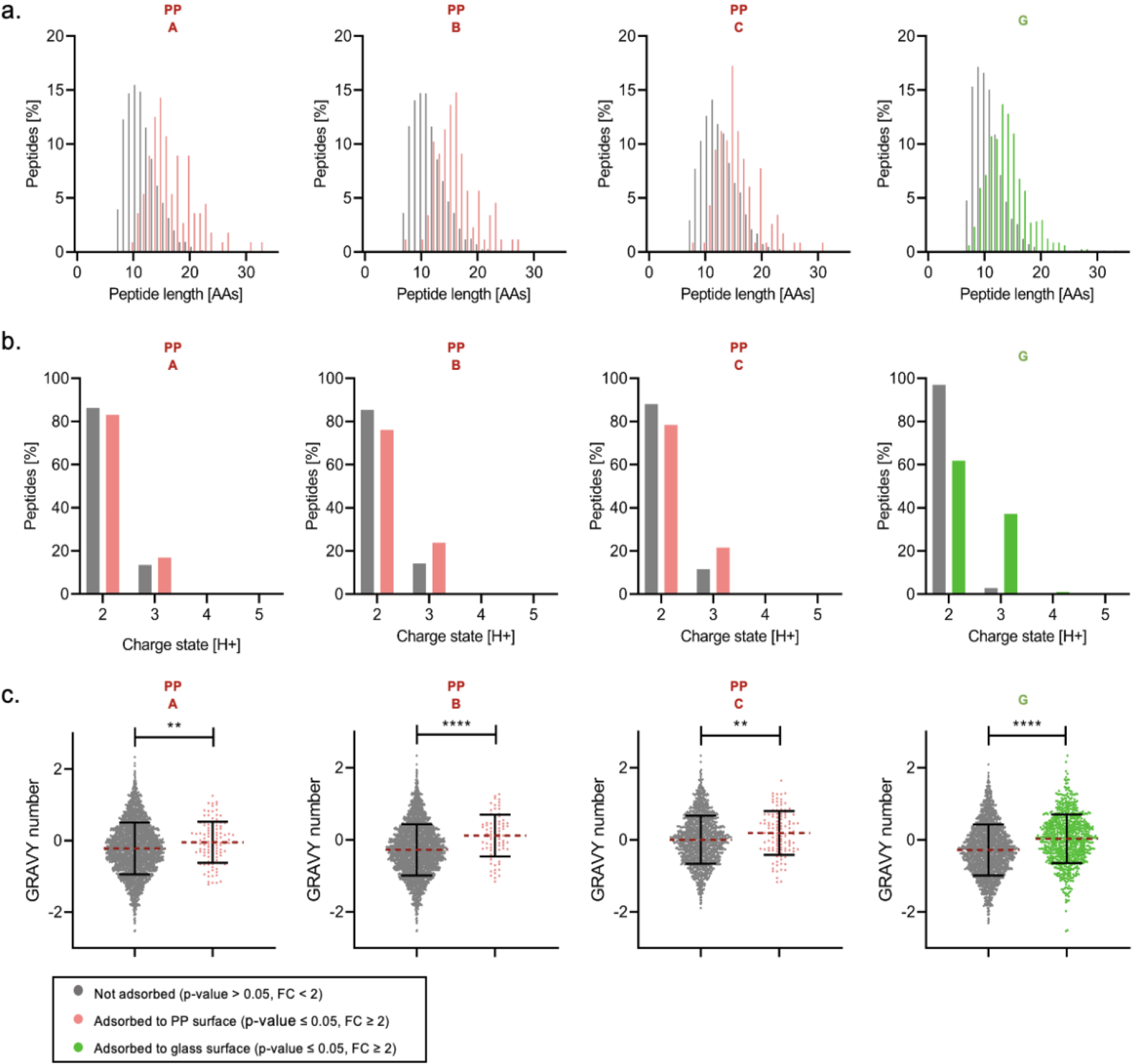
Characteristics of peptides adsorbed to surfaces of polypropylene vials (PP) and of glass vials (G). Analysis of length, charge state and hydrophobicity (based on the hydropathy scale by Kyte and Doolittle)^29^ of adsorbed peptides compared to non-binding peptides in PP vials from manufacturer A, B and C and G vials. Adsorbed peptides are significantly lower abundant after 24h incubation with p-value ≤ 0.05 and ≥ 2-fold-change. a. Peptide lengths of adsorbed and non-binding peptides, b. Charge state comparison between adsorbed and non-binding peptides, c. Gravy number of adsorbed peptides. Significant differences are marked with *: p-value ≤ 0.05, **: p-value < 0.01, ***: p-value < 0.001, ****: p-value < 0.0001, n. s.: p-value > 0.05 in two-sided Wilcoxon rank sum test with continuity correction. APS test results can be found in supplementary Table S3.

We compared the proportions of identified peptide lengths of non-adsorbed peptides to adsorbed peptides (Figure 2. a.) and found that peptides that were non-binding to PP vials had a mean length of 11.2 (A), 11.4 (B) and 12.1 (C) amino acids, while peptides adsorbed to PP vials, showed a significantly higher (>40 %) mean length for all PP-manufacturers (A:17, B:16, C:16). Peptides adsorbed to G, were also longer with a mean of 14 amino acids being adsorbed to G versus 11 amino acid long peptides being non-binding. Comparing the peptide hydrophobicity between adsorbed and non-binding peptides, a significant (p-value < 0.01) higher mean GRAVY-score was observed for peptides adsorbed to PP-vials as well as to G (Figure 2. c.). As a result hydrophobic interaction can be considered as the main reason for peptide adsorption to the hydrophobic polypropylene material^7^. This goes in line with the findings of Kraut et.al. (2009), showing a significant adsorption of hydrophobic peptides to standard PP, investigating a defined peptide mixture, generated from 12 proteins.^17^. Hence, longer peptides are more likely to adsorb to the hydrophobic surface of unmodified PP as they have a higher probability to contain multiple hydrophobic amino acids.

For peptides adsorbed to G vials a significant higher amount of +3 charged peptides (+34.4 %) was observed in comparison to non-binding peptides to G, showing that a higher charge is increasing the adsorption to the glass surfaces. This can be traced back to the higher occurrence probability of positively charged histidine residues at pH 2.7 in longer tryptic peptides. Type 1 borosilicate glass, used for the manufacturing of respective vials, mainly consists of silicon dioxide (SiO_2_).^6^ The oxygen in silica glass has a higher electronegativity than the silicon and a dipole moment arises with partial negative charging of the oxygen. The pI of borosilicate glass is at pH-values ranging from 2-3 ^30^. Consequently, a strong electrostatic interaction via the negative charge of glass surfaces can be expected, leading to the adsorption of positively charged analytes like peptides.

Unexpectedly, also hydrophobic peptides, indicated by a higher GRAVY number of adsorbed peptides, adsorbed to glass surfaces although glass is hydrophilic^32^. Here, a clear differentiation based on chemical characteristics was observable (Figure S3). Adsorbed peptides that were highly charged (> +2) were at the same time less hydrophobic than adsorbed peptides with only +2 charge, dividing the adsorbed peptides into two groups: a highly charged and a highly hydrophobic group. We therefore assume a secondary binging effect of highly hydrophobic peptides to, already adsorbed, highly charged peptides being responsible for the enormous adsorption to G vials.

To gain a deeper understanding of factors underlying the adsorption of peptides to PP and G, we analyzed the amino acid (AA) composition of adsorbed peptides (Figure 3). To analyze the influence of the chemical properties of amino acid on adsorption independently of the peptide-length, only peptides with a sequence comprising 15 amino acids, being the most strongly adsorbed peptides as shown before (Figure 3), were analyzed (Figure 3. a.).

**Figure 3.**
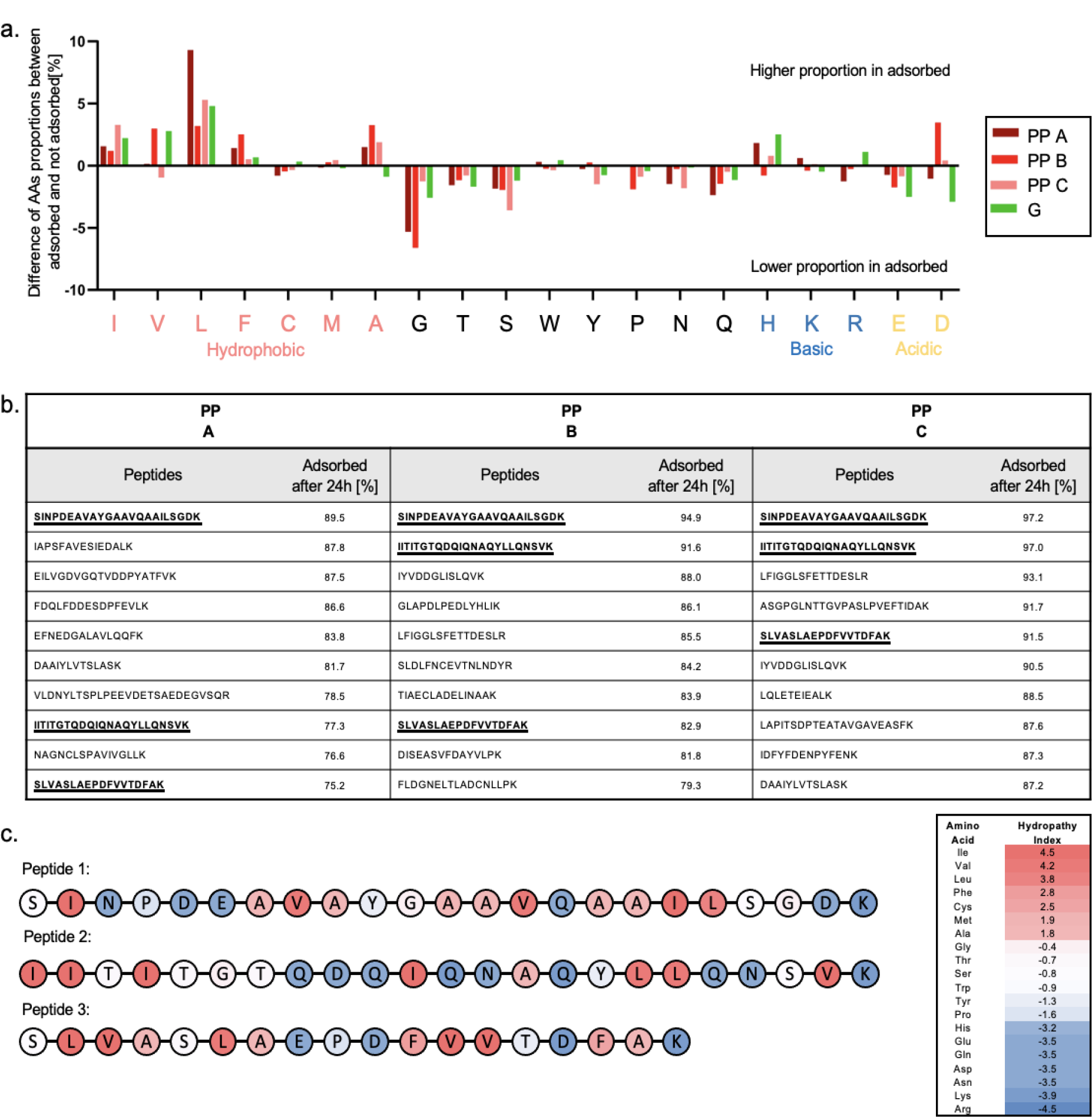
Amino acid composition and hydrophobic distribution of adsorbed peptides. a. Comparison of peptide length (in AAs) of adsorbed and non-binding peptides in polypropylene vials (PP) and glass vials (G) after 24h incubation. Values are calculated based on the amino acid proportion within 15 AAs long peptides and percentage in adsorbed peptides was subtracted from non-binding peptides. Results can be found in supplementary Table S4. b. Top-10 adsorbed peptides in PP vials from manufacturer A, B and C. Peptides present within the top-10 across all 3 manufacturers are marked, c. Hydrophobicity analysis of the 3 top-10 adsorbed peptides in PP vials based on the hydropathy index^29^. The hydropathy index is based on predicted amino acid hydrophobicity in a protein sequence and therefore assumes disulphide bridges between cysteines.^31^ HeLa protein digest reference sample contains alkylated cysteines preventing formation of disulphide bridges and the amino acid can therefore be seen as less hydrophobic in its alkylated form.

Correspondingly with the higher GRAVY number of adsorbed peptides to PP, we found that the hydrophobic AAs alanine, valine, leucine, isoleucine, and phenylalanine were overrepresented in adsorbed peptides, compared to non-binding peptides. Cysteine residues are carbamidomethylated following incubation with iodoacetamide.^33^ As a result, hydrophobic properties of the AA differ from the hydropathy index. Prior analysis, peptides containing oxidized methionine were removed, leading to a reduced sample size for comparing adsorbed and non-adsorbed peptides and low statistical power. As a result, changes in the proportion of cysteine and methionine were not detected. In contrast, the neutral AAs proline, serine, threonine, tryptophan, asparagine, glutamine, tyrosine, and glycine were less abundant in peptides adsorbed to PP-vials compared to non-binding peptides. Peptides adsorbed to G vials showed similar tendencies. In addition, basic amino acids (histidine, lysine, arginine) were significantly overrepresented in peptides adsorbed to G surfaces, while a significant underrepresentation was found for acidic amino acids (glutamate, aspartate). Since the amino acids side chains have pKa values of 3.65 (aspartate) and 4.25 (glutamate), it will predominantly be neutral at the acidic pH of 2.7. However, repulsion of the peptides from the negatively charged borosilicate glass must be considered.

The hydrophobic amino acids are the main drivers of peptide adsorption to unmodified PP surfaces. Interestingly, glycine was underrepresented in adsorbed peptides to PP. This can be linked to changes in the 3D structure of peptides through glycine residues. Especially in the proximity to hydrophobic amino acids, glycine residues induced the formation of ß-sheets.^34^ Induced β-sheet inherit a less linear structure. As a result, smaller numbers of hydrophobic amino acids are exposed to the peptide surface and can interact with the surface material.

Focusing on the top 10 most strongly adsorbed peptides to PP surfaces of all manufacturers, peptide adsorption rates ranging up to 97.2 % of the previous peptide intensity were identified (Figure 3. b.). Three peptides were found among the top 10 adsorbed candidates for vials from all manufacturers and analyzed with respect to their hydrophobicity (Figure 3. c.). The targeted peptides were found to be significantly longer compared to the average of non-binding peptides (Peptide 1: 23 AAs, Peptide 2: 23 AAs, Peptide 3:18 AAs). Additionally, we found that for these peptides, the hydrophobic amino acids were equally distributed across the peptide sequence. The chemical characteristics of frequently adsorbed peptides should be regarded, when selecting vials for experiments focusing on individual peptides. As hydrophobicity was found to be the main driver of peptide adsorption, especially peptides deriving from transmembrane domains are at risk for adsorption to vial surfaces, since they are longer and characterized by high numbers of hydrophobic amino acids.^35^ APS enables a description of adsorption characteristics of a surface even if its exact chemistry is not known.

### The impact of peptide adsorption on protein quantities determined with bottom-up proteomics

The analysis of relative protein abundances for e.g., biomarker discovery, is commonly performed with bottom-up proteomics. Here, proteolytic peptides are identified and quantified by liquid-chromatography tandem mass spectrometry (LC-MS/MS). The sum of obtained peptide abundances for a respective protein is used for its quantification. As a result, the adsorption of individual peptides can significantly alter the calculated protein intensity in a sample. To demonstrate the impact of peptide adsorption on relative protein quantification, protein abundances were compared between control and 24h incubated vials for PP (S, LPB) and glass (S, LR) (Figure 4).

**Figure 4.**
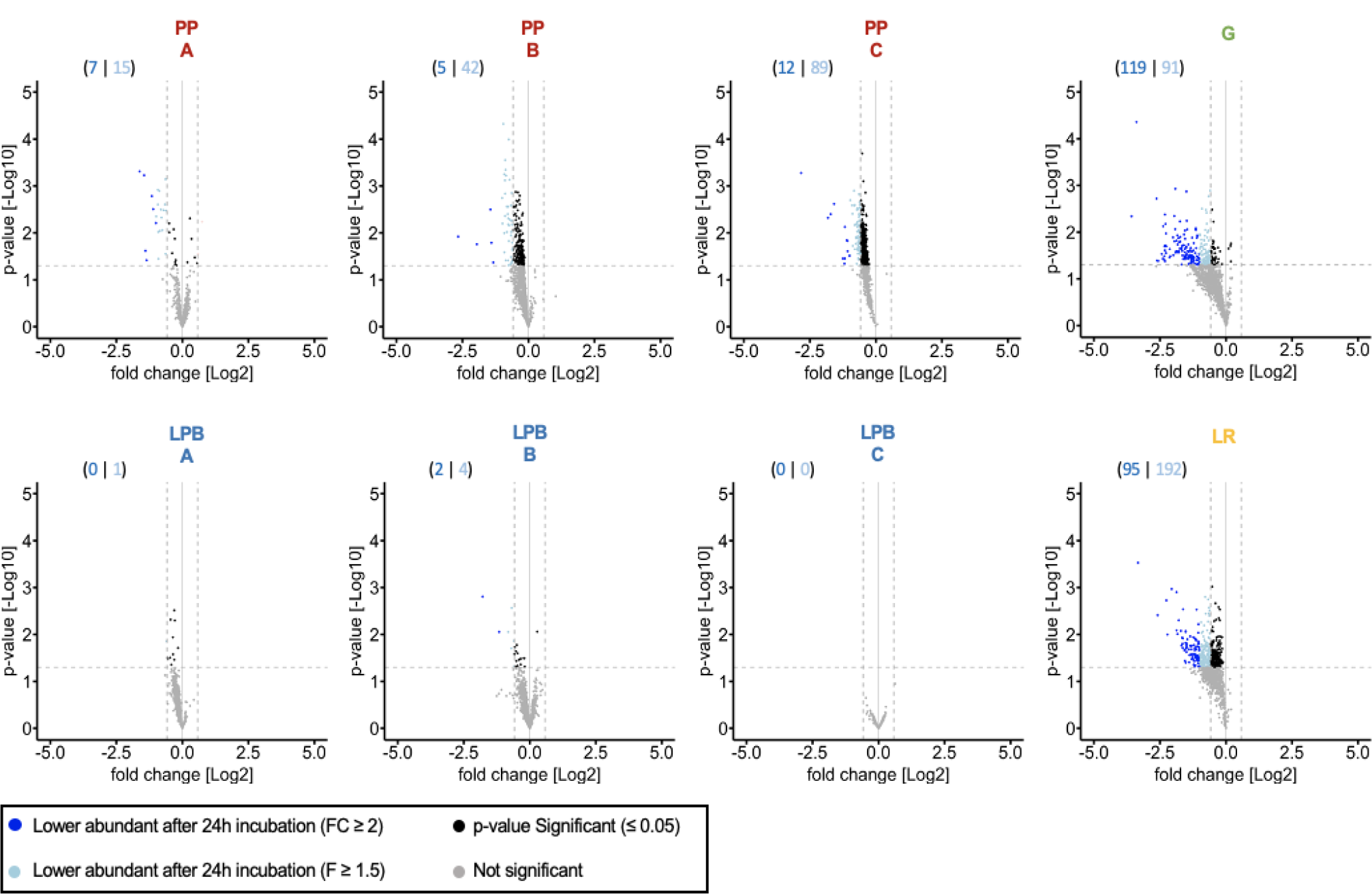
Effect of adsorption of peptides towards surfaces of vials on the quantities of proteins determined with bottom-up proteomics. Volcano plots, plotting the –log10 p-value against the log2 fold change difference for the comparison between the control sample (0h) and the samples incubated 24h, individually determined for low protein binding (LPB)-polypropylene vials, polypropylene (PP) vials, glass (G) vials and low retention (LR) glass vials. A, B and C: Different manufacturers of PP vials. Only proteins being identified across all samples (N=1207) were considered. Quantification was performed based on summed peptide abundances. Proteins identified with a p-value ≤ 0.05 and a fold-change difference ≥ 2 (dark blue) or ≥ 1.5 (light blue) were considered significantly differential abundant. Protein abundances can be found in supplementary Table S6.

In total 1207 Proteins were identified and quantified across all analyzed samples. Welch’s T-testing identified in PP vials 7 (manufacturer A), 5 (manufacturer B) and 15 (manufacturer C) proteins, that showed a p-value significant, two times lower abundance after 24h incubation. Applying a less stringent cut-off, frequently applied in proteomic experiments^36^ and considering proteins that are at least 1.5 times lower abundant after incubation, 15 (manufacturer A), 42 (manufacturer B) and 89 (manufacturer C) additional proteins were identified as significantly changed. For LPB vials of manufacturer C, no abundance difference was detected between control and incubated samples. For manufacturer A, one protein exceeded the 1.5-fold-change cut-off. In accordance with the peptide level, a significant number of proteins was found to be statistically significant lower abundant after 24 hours incubation for LPB vials from manufacturer B (2 proteins ≥ 2fold difference; 4 proteins ≥ 1.5-fold difference).

In G vials, 210 proteins showed a ≥ 1.5 lower abundance and 119 exceeded the ≥ 2-fold-change cut-off. In LR vials a higher number of proteins were found to be significantly adsorbed (≥ 1.5-fold change: 287; ≥ 2-fold change: 95 peptides). Since glass vials are very common for storage of peptide samples in the autosampler directly prior to their injection into the LC-MS/MS system, adsorption can occur. Finally, the storage of samples in PP and G or LR vials can artificially alter protein abundances caused by peptide adsorption and result in false results in differential quantitative proteomics^37^. These results demonstrate that with APS subtle differences between chemically similar surfaces are detectable. Furthermore, APS showed that the surface modification of polypropylene vials was effective in significantly reducing peptides adsorbing to the surface of vials and prevent disturbance of proteomics results. The assay can also be modified and applied for the adsorption analysis in the field of other biomolecules than peptides such as metabolomics^38^ and lipidomics^39^ that are also assessable with quantitative mass spectrometric analysis.

### Automated analysis with APS

To make our APS test applicable for users, an automated script was developed to simplify analysis of adsorption to surfaces. With a simple input of peptides and respective quantitative values of the control samples and samples after incubation, Welch’s T-testing is performed and adsorbed peptides are defined. In the following peptide length, GRAVY as a metric for hydrophobicity based on the Kyte and Doolittle scale^29^ and the charge states of peptides is calculated and visualized comparing “adsorbed” and “not adsorbed” peptide *status* for the tested surface material (Figure S4). An Excel containing all defined peptide metrics is generated.

## CONCLUSION

In this study, we tested the performance and applicability of APS, a new assay for characterizing adsorption-properties of surfaces. APS confirmed that adsorption of molecules to commonly used vial surfaces is a significant phenomenon. With APS the nature of the interactions is characterizable. The choice of material during sample handling is exceptionally important for decreasing the risk of false results. APS is a new tool which will help to find materials as well as parameters minimizing the loss of analytes during sample handling. Furthermore, the significant impact of the loss of peptides due to adsorption on results of bottom-up proteomics studies was demonstrated. Following the findings of this study for conventional bottom-up proteomics workflows, glass vials should be avoided.

## ASSOCIATED CONTENT

### Supporting Information

The Supporting Information is available free of charge on the ACS Publications website.

Supplementary Figures: Figure S1. Principal component analysis and person correlation of incubated polypropylene and glass vials, Figure S2. Overlap of peptides adsorbed to G and LR, Figure S3, Correlation between charge state and GRAVY number of peptides adsorbed to glass vials, Figure S4. APS Test results example of PP tube from manufacturer A. (PDF)

Supplementary Tables: Table S1: Peptide abundances, Table S2: Peptide abundances (log2), Table S3: APS Results, Table S4: AA Occurrence, Table S5: Protein abundances, Table S6: ID (Excel)

### Code availability

The APS test script can be accessed via GitHub under https://github.com/UKE-AGSchlueter/APS_Test.

### Proteomics data availability

The mass spectrometry proteomics data have been deposited to the ProteomeXchange Consortium via the PRIDE^40^ partner repository with the dataset identifier PXD044268. Raw file assignment can be found in supplementary Table S6.

## AUTHOR INFORMATION

### Author Contributions

The manuscript was written through contributions of all authors. / All authors have given approval to the final version of the manuscript.

### Notes

The authors declare the following competing financial interest: This study has been performed in cooperation between the Core Facility Mass Spectrometric Proteomics and Eppendorf SE, the manufacturer of vials tested in this research. D.H. is employee of Eppendorf SE.

## ACKNOWLEDGMENT

This study was supported by grants from the Deutsche Forschungsgemeinschaft (DFG) (INST 337/15-1, INST 337/16-1, INST 152/837-1 and INST 152/947-1 FUGG).

## Notes

### Summary of Updates

Change of last-authorship Correction of conflict of interest statement

