## Supplementary Figures for "Assay for characterizing adsorption-properties of surfaces (APS) used for sample preparation prior to quantitative omics"

Figure S1. Principal component analysis and person correlation of incubated polypropylene and glass vials.

Figure S2. Overlap of peptides adsorbed to G and LR.

Figure S3, Correlation between charge state and GRAVY number of peptides adsorbed to glass vials.

Figure S4. APS Test results example of PP tube from manufacturer A.

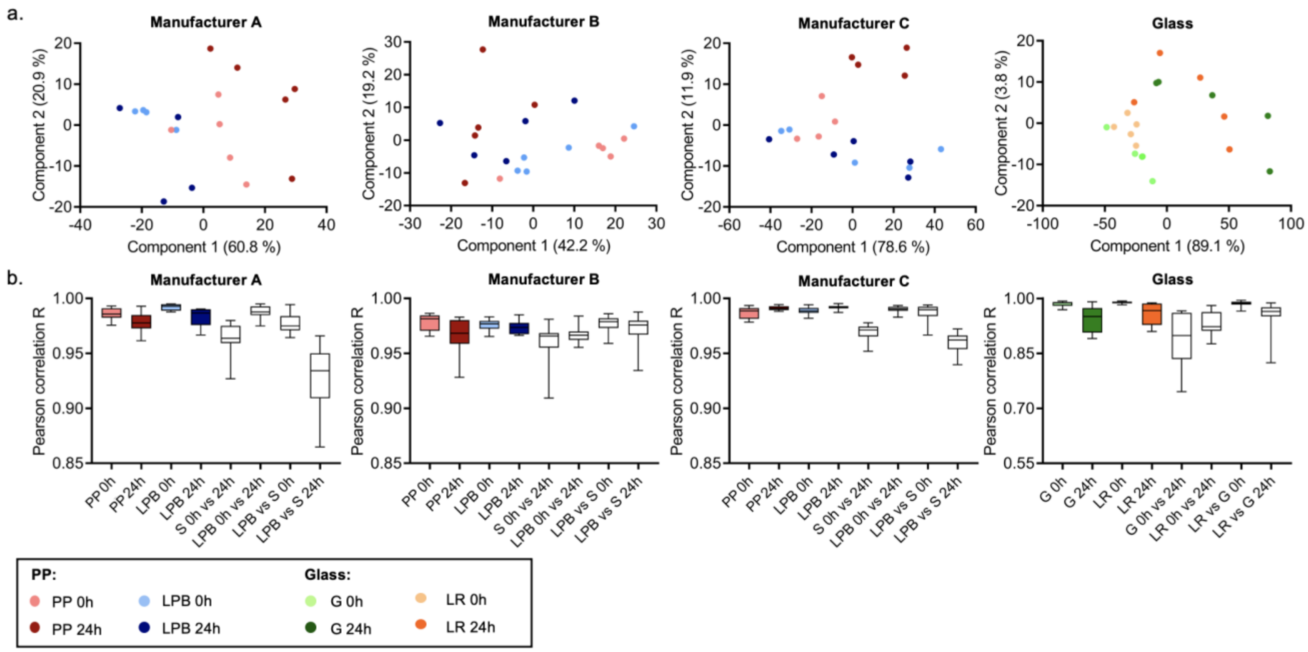


Figure S1. Principal component analysis and person correlation of incubated polypropylene and glass vials. Incubation was performed in polypropylene (PP) and low protein binding (LPB) vials from competitors A, B and C and in glass vials (G) and low retention (LR) glass vials. Measurement of the control sample (0h) and the sample incubated 24h. For each manufacturer and sample (0h sample; 24h sample) n=5 replicates were measured. For further analysis only peptides (N=3531) were used which have been detected across all samples a. Scatter plot visualization of principle components 1 and 2 from the principal component analysis of peptides from the tryptic HeLa peptides control sample (0h) and the samples incubated 24h. Log2 transformed values were used. b. Boxplot visualisation of Pearson correlation coefficients between different vial types and samples for each manufacturer. In boxplots, 50% of the data points are inside the box (Q1 (Quartile 1) being the lower bound of the box (25%), Q3 being the upper bound of the box (75%)). Whiskers show all values beyond the box without outliers. Outliners were defined as Q3 + 1.5 * IQR (Interquartile range) (upper outlier) and Q1-1.5 * IQR (lower outlier). IQR being Q1–Q3.

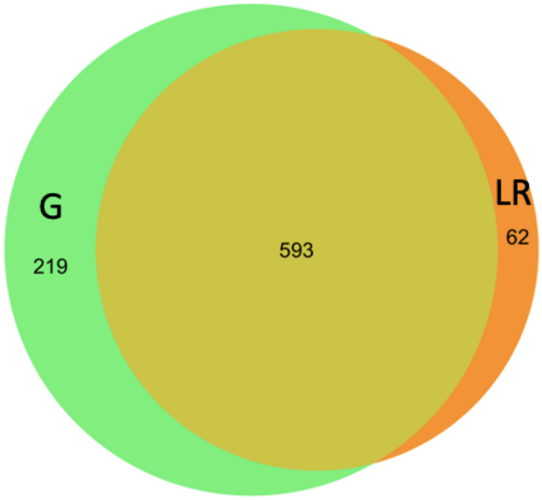


Figure S2. Overlap of peptides adsorbed to G and LR. Venn Diagram representing the overlap of peptides that adsorbed to the standard glass (G) and low retention (LR) glass surface.

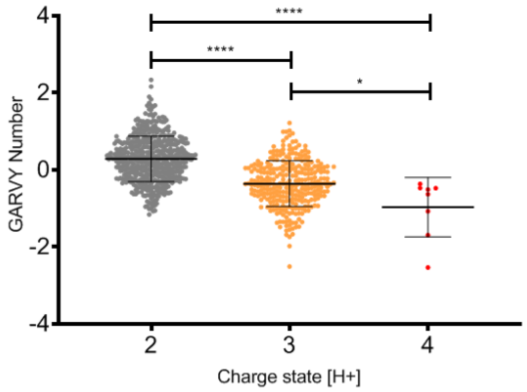


Figure S3. Correlation between charge state and GRAVY number of peptides adsorbed to glass vials. Analysis of hydrophobicity (based on the hydropathy scale by Kyte and Doolittle)[26] of adsorbed peptides compared to non-binding peptides in standard glass vials (G). Adsorbed peptides are significantly lower abundant after 24h incubation with p-value ≤ 0.05 and ≥ 2-fold-change. Non-binding peptides are not p-value significant or fold-change significant. Gravy number of adsorbed peptides in correlation to the occurring charge states +2, +3 and +4 of the peptides. Significant differences are marked with *: p-value ≤ 0.05, **: p-value < 0.01, ***: p-value < 0.001, ****: p-value < 0.0001, n. s.: p-value > 0.05 in two-sided Wilcoxon rank sum test with continuity correction.

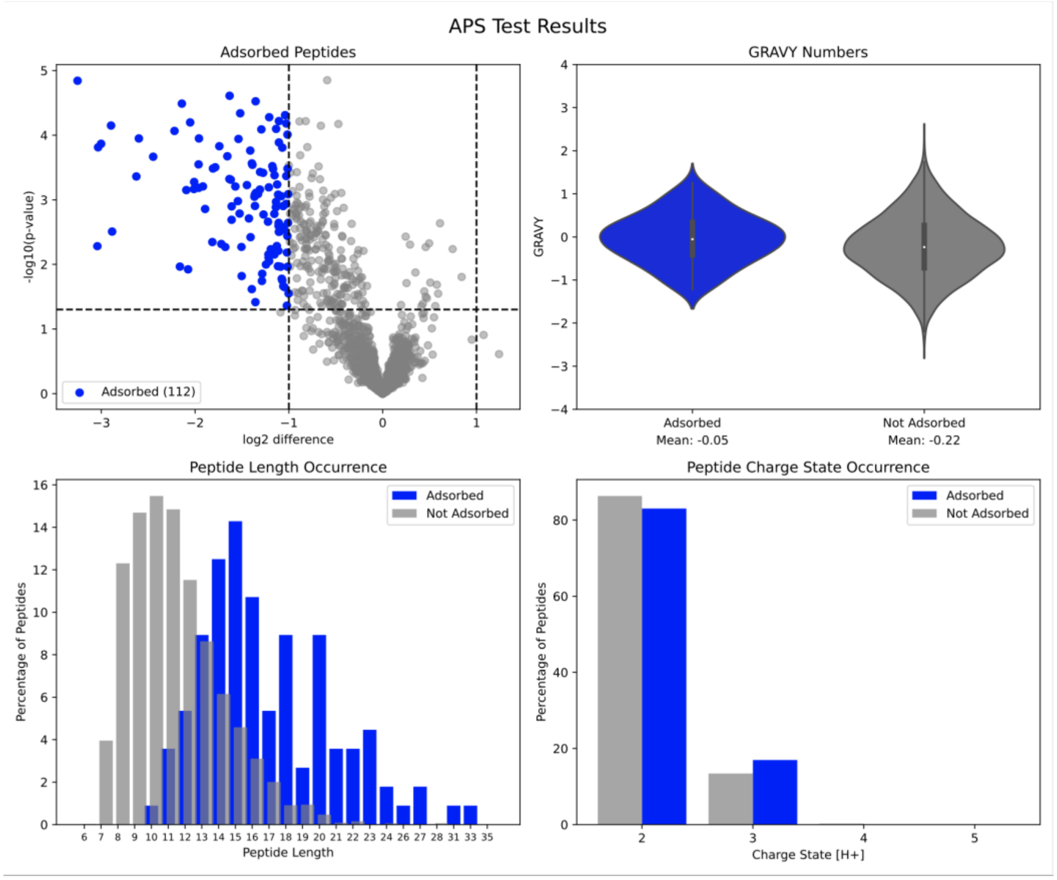


Figure S4: APS Test results example of PP tube from manufacturer A. Upper left: Volcano plot based on Welch's t-test results, Upper right: Violin Plots of GRAVY numbers between adsorbed and not adsorbed peptides, Lower left: Peptide length occurrence in adsorbed and not adsorbed peptides, Lower right: Peptide charge state occurrence for adsorbed and not adsorbed peptides.
